# Comprehensive calculation of product and ATP yields for glucose fermentation

**DOI:** 10.64898/2026.10.12.758529

**Authors:** Maaike Remeijer, Christoff Odendaal, Rebeca Gonzalez-Cabaleiro, Frank Bruggeman

**Affiliations:** Systems Biology Lab, A-LIFE, AIMMS, Vrije Universiteit Amsterdam; Environmental Biotechnology Section, Department of Biotechnology, Delft University of Technology, the Netherlands

## Abstract

Lignocellulose sugars are degraded anaerobically by microbial communities as part of the carbon cycle, yielding short- and medium-chain fatty acids as intermediates and carbon dioxide and methane as final products. The underlying metabolic network is branched and its product yields depend on environmental conditions. We compiled a pan-catabolic network of the anaerobic digestion of glucose into 11 fermentation products, including alcohols, acids, and ketones. It consists of 75 reactions and incorporates energy conservation via substrate-level phosphorylation and transport-coupled phosphorylation, with ferredoxin, NADH, and quinol as electron carriers. We calculated all net conversions of glucose into fermentation products and their ATP yields. We find 195,629 unique conversions (about 50% with non-zero ATP yield), including known and many novel ones. Conversions with identical product stoichiometry can differ in ATP yield: glucose to propionate and acetate is achieved by different subnetworks with 23 distinct ATP/glucose yields, and even glucose to lactate shows 12 yields ranging from 0 to 4.5. The driving force of the conversions decreases with ATP yield, whereas the number of reactions used does not correlate with it. The known catabolic conversions underlying anaerobic digestion might thus be only the tip of the iceberg.

## 1 Introduction

About 90% of plant biomass is lignocellulose, making it the most abundant renewable energy source on Earth (Sulis et al., 2025). It amounts to 57% of Earth’s biogenic carbon (Bar-On et al., 2018). Lignocellulose is composed of cellulose (30-50%; a glucose polymer), hemicellulose (20-30%; contains glucose and mannose), and lignin (5-25%; polyphenols). Its components are all degraded by microbes, as part of the carbon cycle. Glucose is the main component of lignocellulose, its breakdown by microbial communities is one of the most active processes on Earth. This is mostly done anaerobically by microbial communities. It occurs in soils, guts of many species, and fermented foods (sauerkraut, wine, kefir, etc.). Those digestions are sometimes incomplete, ending at an intermediate. In applications, such as product formation from wastewater streams, steering anaerobic digestions towards desired intermediate fatty acids is a process-engineering goal (Khan et al., 2016).

Nearly all carbon products formed during glucose fermentation are byproducts of catabolism which synthesizes charged energy carriers (i.e., ATP) to fuel anabolism (Spormann, 2023). In anaerobic digestions, these carbohydrate products of fermentation can subsequently act as the energy source of methanogenic archaea, which produce carbon dioxide and methane as final products (Buan, 2018; Gujer & Zehnder, 1983). The sequential nature of anaerobic digestion (acidogenesis *→* metanogenesis) is, in part, governed by thermodynamics: fermentation yields more energy per catalytic step than the full digestion of glucose to carbon dioxide and methane (González-Cabaleiro et al., 2015). This effectively creates two niches, one for fast growing fermenters and another for slower-growing methanogens.

The product profile and rate of a fermentation depend on the precise conditions. Changing environmental conditions (pH, temperature, salinity, gas phase composition, etc.) can lead to very different product profiles arising from fermentations – even of the same microbe/community growing on the same substrate (Bühlmann et al., 2022; Dabrock et al., 1992; Temudo et al., 2007). Humans from all over the world have, for thousands of years, been tacitly applying this knowledge in designing selective conditions to enhance the safety, longevity, sensory, and functional properties of fermented foods, from kimchi to wine (Auchtung et al., 2025).

The catabolic networks of specific microbial species are often characterized by their net conversion. An example of such a net conversion is glucose + 2 ADP + 2 Pi *→* 2 lactate + 2 ATP for lactate fermentation or glucose+ 3 ADP+ 3 Pi *→* 1.33 propionate+ 0.67 acetate+ 0.67 CO_2_ + 3 ATP for mixed propionate and acetate fermentation. Such net conversions can be calculated from the stoichiometry of the associated catabolic reactions. Besides the pathway stoichiometry, two more constraints determine the feasible net conversions that can arise from a network (Bruggeman et al., 2026): 1) the requirement for steady-state balancing of the production and consumption rates of all redox carriers and pathway intermediates; 2) reaction reversibility (physiologically or thermodynamically), which determines which reactions are reversible. Net conversions can be calculated from a stoichometric model of a metabolic network by enumerating all its elementary flux modes (EFMs) and determining their net conversions (Kamp & Schuster, 2006; Schuster et al., 1999, 2000). Nowadays, several packages exist for this purpose (e.g. EFMTool, Terzer and Stelling (2008)).

EFMs are steady-state flux routes of metabolic networks that have no redundant reactions (making them minimal) and agree with reaction irreversibility constraints. EFMs are considered elementary, because any steady-state flux distribution of a metabolic network can be written as a linear combination (with positive coefficients) of its EFMs. A feature of an EFM, due to its minimality, is that it has one degree of flux-freedom (i.e., one independent flux). Accordingly, assigning a flux value to one of its reactions suffices to determine all of its other flux values by balancing the synthesis and degradation fluxes of all the metabolic intermediates of the EFM. Researchers focused on microbial metabolism have historically been interested in ATP or carbon byproduct yield per unit of substrate consumed. For this reason, they conventionally constructed catabolic pathways by setting the flux of the first reaction in the pathway (typically the substrate import step) to a fixed value (typically 1). Because of this, famous textbook catabolic pathways are often elementary flux modes (Schuster et al., 2000); for example, this is the case for the pathways drawn in by Spormann (2023). Because the flux routes with maximal product yield on a substrate are always elementary flux modes, EFMs are also the goals of metabolic engineering problems (Klamt et al., 2025) and the outcome of flux balance analysis when one flux constraint in the (genome-scale) stoichiometric model is active (Gagneur & Klamt, 2004).

Each EFM leads to a single net conversion (Bruggeman et al., 2026; Schuster et al., 2000). Since different EFMs can correspond to the same net conversion, (many) more EFMs tend to exist than net conversions. Enumerating net conversions without first enumerating EFMs first, however, is not (yet) possible. Net conversions can only be computed from all EFMs. (Elementary conversion modes (Clement et al., 2021) are a different mathematical object). Computation of all net conversions of a genome-scale stoichiometric model is beyond current computational capabilities, due to an understood combinatorial explosion of EFMs (Kelk et al., 2012). Enumeration of all EFMs and their net conversions is, however, feasible for medium-sized metabolic networks such as the pan-catabolic network considered in this paper.

The aim of this work is to compute the stoichiometric diversity of the fermentative net-conversions of glucose into acids, alcohols, and ketones. We assembled a comprehensive pan-species network of the key reactions underlying glucose fermentation for ATP synthesis. We calculated all its EFMs and net conversions, which we compare on the basis of their product stoichiometries, ATP yield, and Gibbs free energy change at (metabolic) standard conditions. We find many more net conversions than have been described in literature. We discover that net conversions can exist that have the same glucose to product stoichiometries, but differ in their ATP yield on glucose. That we find so many net conversions may either imply that experimentalists have so far only seen the tip of the iceberg or that physiological constraints or trade-offs exist that prevent particular net conversions from occurring in nature.

## 2 Results

### 2.1 Pan-catabolic network of anaerobic fermentation of glucose

We compiled a pan-catabolic reaction network of typical catabolic pathways from different microorganisms that ferment glucose anaerobically (Figure 1). The network consists of glycolysis (Embden-Meyerhof-Parnas pathway), the non-oxidative pentose phosphate pathway, the reductive branch of the TCA cycle, and several pathways to common fermentation products. This includes proton-motive force (PMF)-producing reactions (proton-pumping), hydrogenases, and ATP synthase.

**Figure 1:**
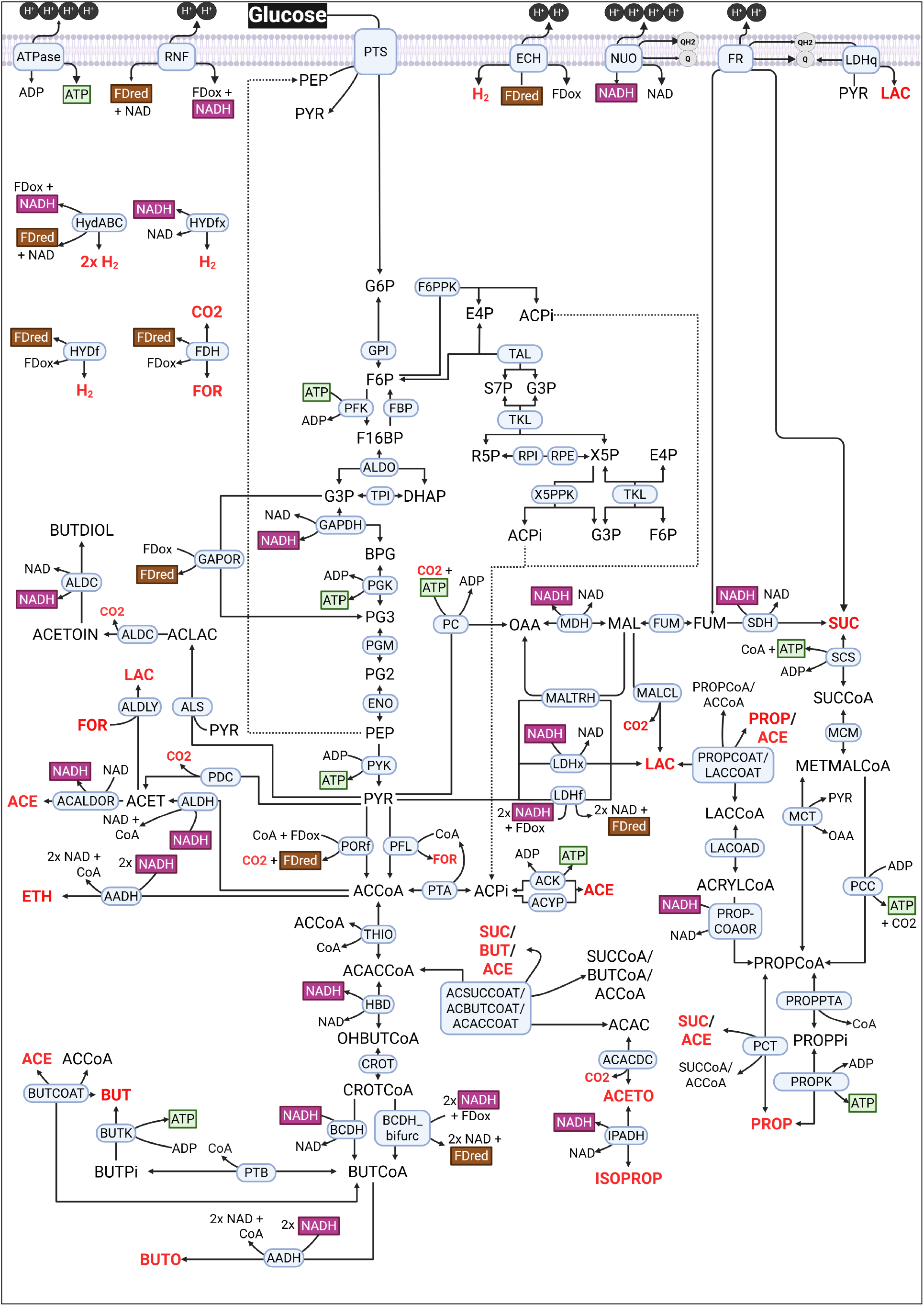
The pan-catabolic network of anaerobic fermentation of glucose. The shown reactions were compiled from literature and are all associated with glucose fermentation pathways that produce the following carbohydrate products (indicated in red and boldface): acetate (ACE), butyrate (BUT), propionate (PROP), succinate (SUC), ethanol (ETH), butanol (BUT), lactate (LAC), formate (FOR), acetone (ACETO), isopropanol (ISOPROP), and butane-2,3-diol (BUTDIOL). The off-gases H_2_ and CO_2_ are also considered products. Not indicated are inorganic P_i_, H_2_O, and H^+^. The stoichiometric model of this network considers all reactants of each reaction, ensuring that all reactions are chemical-elemental balanced.

For the aforementioned fermentation pathways, the point of departure was the chapter on *Fermentative Metabolism* by Spormann (2023). In this chapter, pathways are described for the fermentation of glucose into lactate, acetate, butyrate, ethanol, propionate, acetone, butanol, formate, and butane-2,3-diol. We expanded the network to include two other (industrially relevant) fermentation products: succinate (Song & Lee, 2006) and isopropanol (dos Santos Vieira et al., 2019). In addition to including the pathways described above, we searched the literature, including the KEGG (Kanehisa & Goto, 2000; Kanehisa et al., 2025) and UniProt databases (“UniProt: the universal protein knowledgebase in 2025”, 2025), and added additional reactions. A detailed network description is provided in **Supplementary Text S1**.

ATP can be produced in the network via a number of substrate-level phosphorylation (SLP) reactions, or through transport-coupled phosphorylation (TCP) (Spormann, 2023). Although the role of the proton motive force (PMF) in ATP production has received much attention recently (Lane, 2017), it is known that microbes also harness PMF for the co-transport of nutrients across their cellular membranes (Konings et al., 1981) or dissipate it unproductively (Baronofsky et al., 1984). In our pan-catabolic model, transport has been simplified to uniport only and no proton leak is included, which means that PMF is fully harvested for ATP production via ATP synthase, with a stoichiometry of 4 H^+^ per ATP, as reported by Buckel and Thauer (2013) and Schoelmerich et al. (2020).

We only consider NADH in the network and not NADPH. We did, for instance, not consider a NADPH-dependent GAPN nor the oxidative pentose phosphate pathway. This was because of our intention to use this network to calculate net steady-state conversions of glucose into (excreted) fermentation products and into ATP (fueling anabolism). And, although NADPH is also used to fuel anabolism in many microbes, the exact NADPH/ATP stoichiometry per unit biomass is likely condition- and species-dependent. All our calculations can be repeated with a preset anabolic of NADPH/ATP stoichiometry, we provide all the information to do this. This would then require adding the following reaction *ATP* + *x NADPH → anabolism* to the network and the consideration of ATP and NADPH as internal metabolites in the stoichiometric model.

Although not exhaustive, the resulting network is an extensive representation of the catabolic capabilities present among fermentative anaerobes. A detailed network description with relevant literature citations is provided in **Supplementary Text S1**. This information is also in an Excel file (Supplementary File 1), which summarizes three inputs required for converting the network to a stoichiometric model: i. each reaction written in a elementally balanced format, ii. metabolites assigned as either internal or external, and iii. reactions defined as either reversible or irreversible.

The difference between internal and external metabolites is that the net synthesis and consumption rates of internal metabolites are balanced at steady-state conditions for the network, while the external metabolites can display a net flow in or out of the network at steady state. Hence, only the external metabolites end up in the net conversion of the network. We considered the following external metabolites: glucose, water (H_2_O), non-pumped protons (H^+^), inorganic phosphate (Pi), carbon dioxide (CO_2_), hydrogen (H_2_), ATP, ADP, and the 11 fermentation products. The pumped protons constituting the PMF, which can be harvested for ATP production (referred to in the model as ‘IONpumped’), and all electron carriers are treated as internal metabolites and their respective fluxes balance at steady state.

The stoichiometric model we constructed considered 62 internal metabolites, 19 external metabolites and 141 reactions, including 75 conversions and 38 transport and exchange reactions. Any self-consistent description of the stoichiometric model gives rise to net conversions that are elementally balanced (which we confirmed).

### 2.2 Enumeration of all net conversions of glucose into fermentation products

We used EFMTool (Terzer & Stelling, 2008) to enumerate all the elementary flux modes (and their associated net conversion) of the stoichiometric model of the pan-catabolic network. EFMs are each steady-state flux vectors with all fluxes running in their feasible directions and the associated reaction network has no redundant reactions. The rates of reactions involving external metabolites indicate their in- or outflow rate of the network and suffice to determine the net conversion (Schuster et al., 1999, 2000). We normalized all net conversions per 1 mol glucose consumed.

EFMTool enumerated 60,259,323 EFMs. Many of them gave rise to the same net conversion, we ended up with 195,629 different net conversions. These involved 110,895 unique product stoichiometies, irrespective of their ATP stoichiometry. We associated each net conversion with a single EFM: the EFM involving the minimal sum of fluxes (normalized for the uptake of 1 glucose). In the entire set of net conversions all products were indeed made (Figure 2A). (This figure does not indicate which combinations of fermentation products are formed.) Acetate proved the most common fermentation product and ethanol the least common. Approximately half of the net conversions exhibited a net production of ATP (Figure 2). This left 105,286 net conversions that have a non-zero ATP yield on glucose.

**Figure 2:**
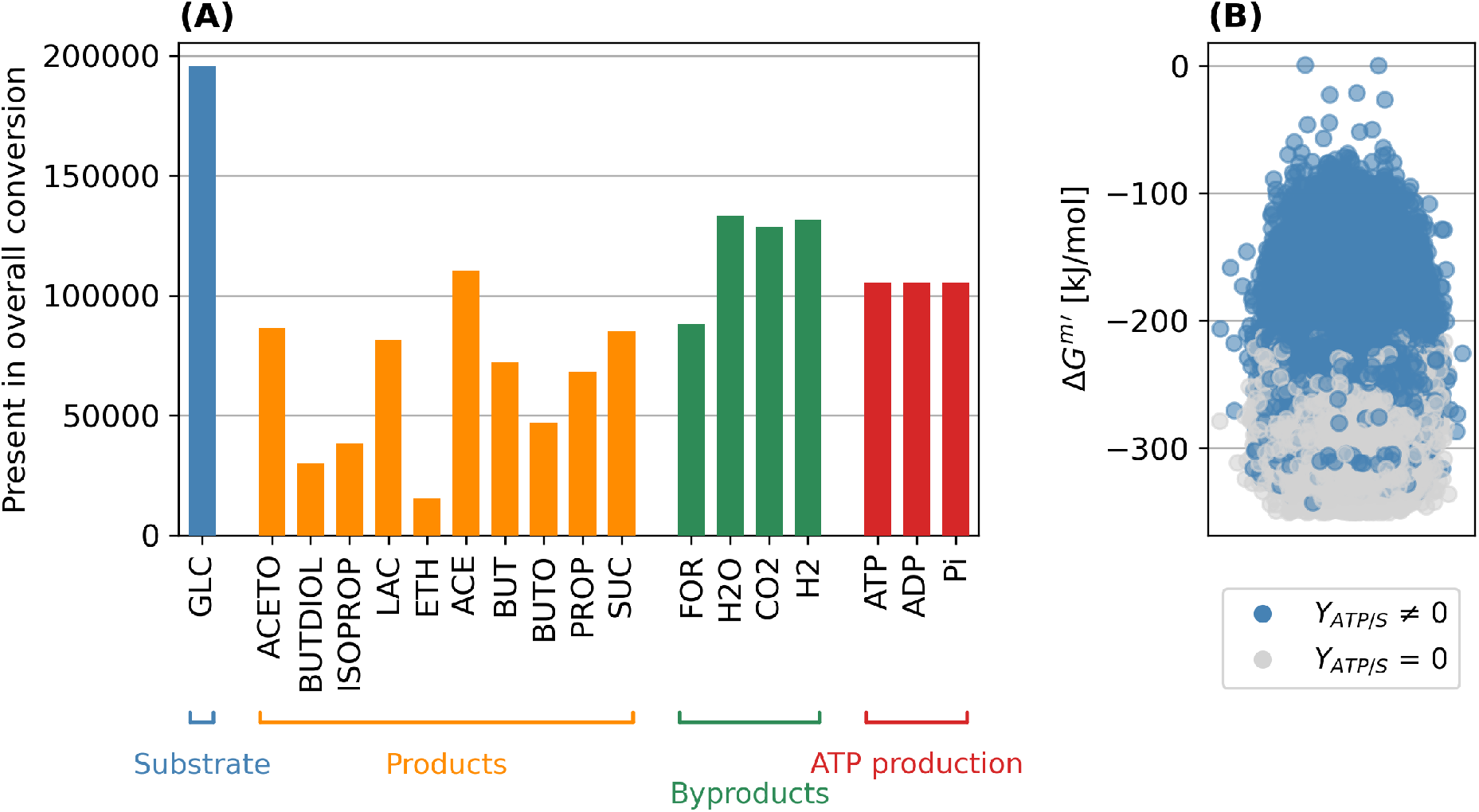
General statistics about the enumerated EFMs. (A). The number of EFMs in which each external metabolite participates, categorised by role: substrate (blue), product (green), or ATP synthesis from ADP and P_i_ (orange). Abbreviations: GLC, glucose; ACETO, acetone; BUTDIOL, butanediol; ISOPROP, isopropanol; LAC, lactate; ETH, ethanol; FOR, formate; ACE, acetate; BUT, butyrate; BUTO, butanol; PROP, propionate; SUC, succinate. Note that the non-pumped protons do not appear. This is due because we use protonated versions of the reactants (KEGG convention) such that protons are made reduced electron carriers are made and those are then balanced during electrion carrier oxidation. (B) The physiological Δ*G* (Δ*G*^*m′*^, pH 7, 1 mM for all metabolites) in kJ/mol glucose of each overall stoichiometry. All net conversions are shown as a dot in this figure. Those that do not yield ATP (i.e. *Y*_*AT P/X*_ = 0) are shown in grey and tend to have more negative Δ*G*^*m′*^ than those that do make ATP (i.e., *Y*_*AT P/X*_*≠* 0; shown in blue).

Figure 2B indicates the Gibbs free energy change of all net conversions (at metabolic standard conditions, Δ*G*^*m′*^). Those that do not make ATP (*Y*_*AT P/S*_ = 0) have the most negative Δ*G*^*m′*^ and, therefore, dissipate the most free energy.

### 2.3 Different metabolic pathways with the same glucose into catabolic products conversion but with different ATP yields

The set of 105,286 net conversions that produce ATP corresponds to 45,392 net conversions with different stoichiometries of fermentation products. On average, 1.76 net conversions occur that have the fixed product stoichiometry but differ in their ATP yield (the median is 1.0). The product stoichiometry with the highest number of unique ATP yields is the production of acetone and has 97 distinct ATP yield. An overview of this is shown in **Supplementary Figure S2**.

An example of a conversion of glucose into catabolic products that occurs with different ATP yields is shown in Figure 3. These examples consider fermentation of 1 glucose to 4/3 propionate, 2/3 acetate and 2/3 CO_2_. This is typical of propionic acid bacteria such as *Propionibacterium freudenreichii* and is, for instance, important in cheese production (Bücher et al., 2021). The three examples displayed in Figure 3 correspond to the following ATP yields: one with a low ATP yield (1.56 ATP/Gluc), one with a moderate (2 ATP/Gluc), and one with a high yield (5 ATP/Gluc). The striking range of possible ATP yields for the same overall conversion arises from the variety of combinations of reactions that all obey steady state. Note that these EFMs all have product yields corresponding to known microbial behaviours, but none follow a known pathway stoichiometry (cf. Spormann (2023)).

**Figure 3:**
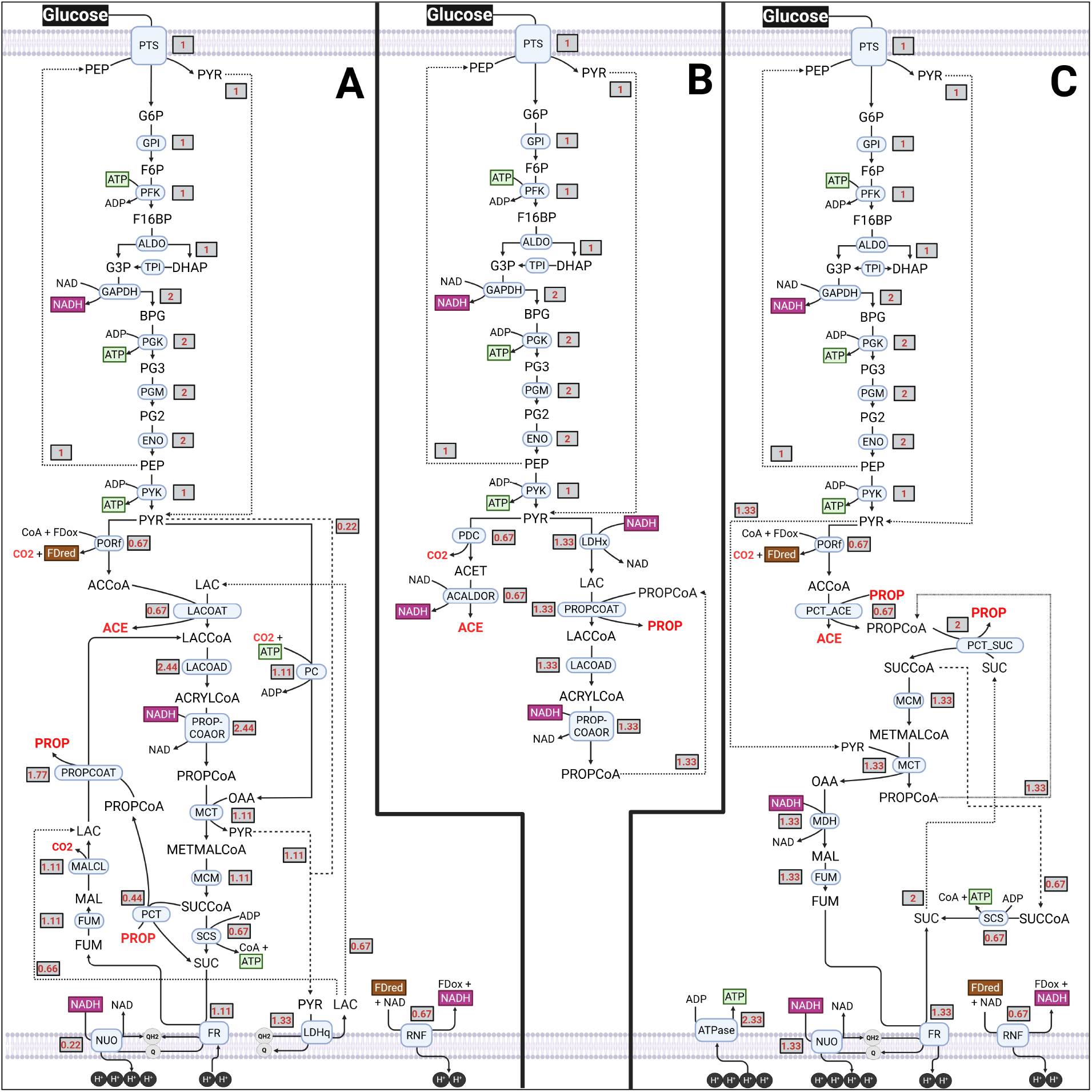
Three propionate-acetate fermentations with different ATP yields. Three EFMs with identical yields of propionate (1.33 Prop/Gluc) and acetate (0.67 Ace/Gluc), but different yields of ATP, are compared: low yield, 1.56 ATP/Gluc (**A**); moderate yield, 2 ATP/Gluc (**B**); and high yield, 5 ATP/Gluc (**C**). Final products in boldface and red, fluxes in grey boxes.

The lowest ATP yield (Fig. 3A) involved the pathway with the most reactions (i.e., 27 reactions, **Table S2**). It relies on a convoluted pathway to produce pyruvate, which is carboxylated to oxaloacetate, which dissipates ATP (Fig. 3A). It contains two PMF-forming steps (i.e., NADH-quinone oxidoreductase, NUO, and the RNF) and one PMF-dissipating step (i.e., fumarate reductase, FR). FR and NUO are coupled via the quinol/quinone (QH_2_/Q) redox pair, which is in turn coupled to a membrane-bound, QH_2_/Q-linked lactate dehydrogenase (LDHq).

The shortest pathway was the EFM with a moderate ATP yield (2 ATP/Gluc, Fig. 3B). It consisted of 17 reactions and lacks proton pumps. It contains reactions that are more typical of the textbook understanding of propionate/acetate producers.

To reach a yield of 5 ATP/Gluc, the third stoichiometry relied on a large network (23 reactions, Fig. 3C) with three PMF-forming reactions (NUO, FR, RNF). The PMF was harvested for ATP production, via an ATP synthase, producing 2.33 ATP on top of the 2.67 produced via substrate-level phosphorylation.

### 2.4 Different metabolic pathways with the same net conversion

Collapsing the list of enumerated EFMs (60 million) to contain only the shortest route for each conversion (195,629 EFMs) was a strategy to make the dataset more manageable. However, it masks an even greater potential diversity. To give an impression of what is being neglected in this process, Fig. 4 shows three EFMs with identical overall conversions, including identical ATP yields:

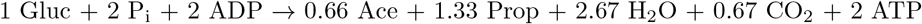

**Figure 4:**
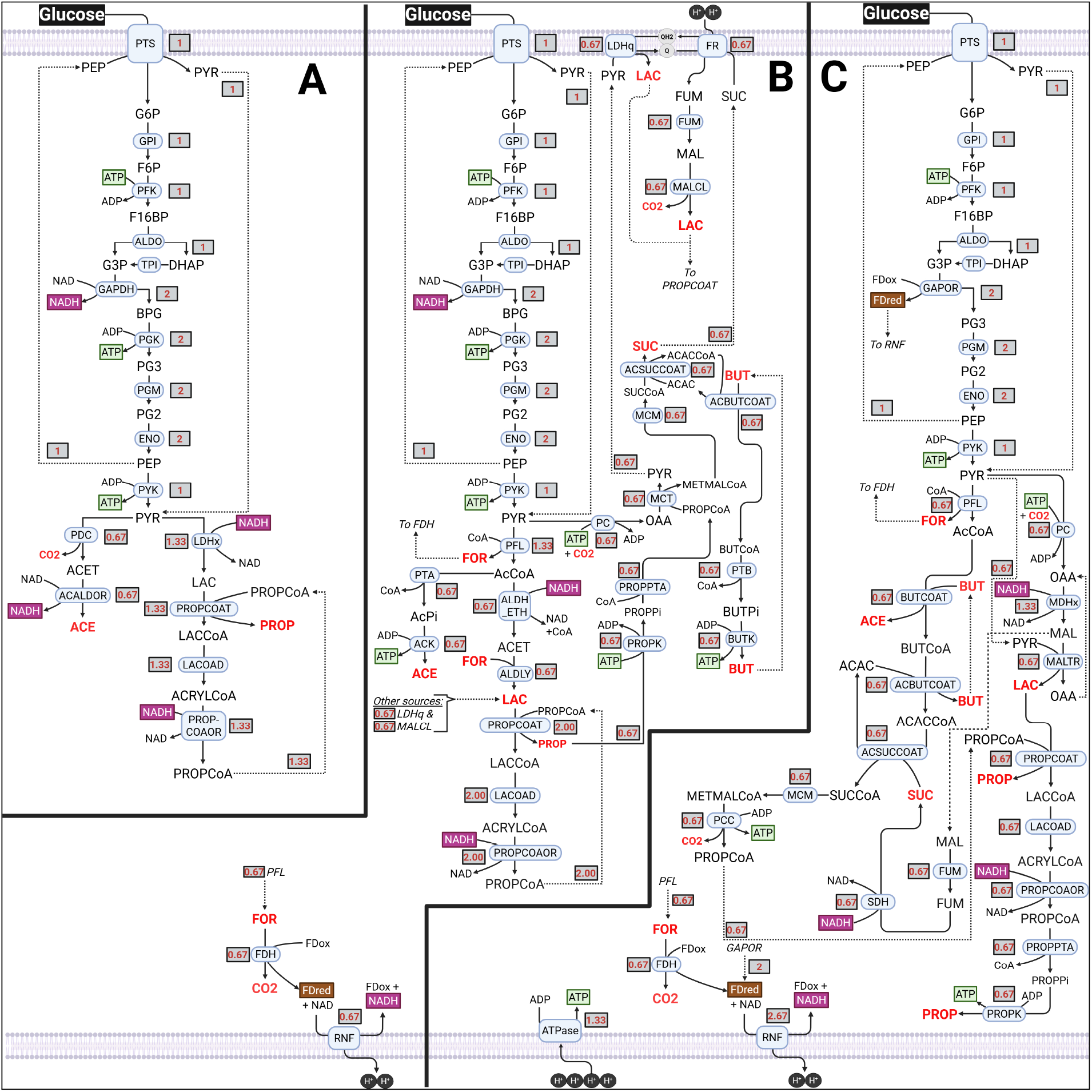
Three propionate-acetate fermentations with identical overall conversions. Three EFMs with identical yields of propionate (1.33 Prop/Gluc) and acetate (0.67 Ace/Gluc) and ATP (2 ATP/glucose) are compared: the shortest, 17 reactions (**A**); the longest, 35 reactions (**B**); one containing the least common reaction for this particular conversion - propionyl-CoA carboxylase, PCC (**C**). Final products in boldface and red, fluxes in grey boxes. Note that the overall conversions performed by these EFMs are identical in every sense, including P_i_ consumption and H_2_O and CO_2_ production.

We show the shortest (Fig. 4a, 17 reactions; identical to Fig. 3b), the longest (Fig. 4b, 35 reactions), and an EFM containing the reaction which is least frequently active in the set of EFMs underlying the above-mentioned overall conversion: propionyl-CoA carboxylase (PCC; Fig. 4c). The EFMs can be found in **Supplementary Table S3**. Fig. 4b shows, again, how coupling proton pumps can lead to a longer EFM: the production of PMF by RNF is harvested by FR to convert succinate to fumarate. This then also yields quinol, which acts as an electron donor for the conversion of pyruvate to lactate, which can, in turn, be converted to propionate further downstream. CoA transferases also allow non-intuitive routes to arise. For example, butyrate is internally recycled in Fig. 4b as CoA acceptor from acetoacetyl-CoA, yielding an ATP-producing cycle. Fig. 3c also makes use of butyrate as CoA carrier. Notably, this pathway relies on a ferredoxin-coupled glyceraldehyde-3-phosphate dehydrogenase (GAPOR; see Spormann, 2023). The reduced ferredoxin yielded by this reaction is then used directly by RNF to produce PMF for ATP synthesis.

In summary, sketching different EFMs that can lead to the same overall conversion with different (Fig. 3) or even the same ATP yield (Fig. 4) shows how cycling and redox-coupling might allow for a vast diversity of underlying pathways. While it is, for instance, often assumed that PMF is intended for ATP production or substrate import (as in all the examples discussed by Spormann (2023)), we show that PMF might drive the reduction of an internal substrate, e.g. via FR. Moreover, pathways with no net butyrate production might contain butyrate as CoA-transferring intermediate, a possibility never addressed in any of the literature that we have consulted.

### 2.5 Performance comparison of net conversions

A feasible net conversion satisfies two criteria: it produces a net amount of ATP (considered above) and is thermodynamically feasible (a negative Δ*G*). We calculated the Gibbs free energy under standard physiological conditions (Δ*G*^*m′*^; 1 mM concentrations, pH 7) for each overall stoichiometry using eQuilibrator (Noor et al., 2013). The resulting Δ*G*^*m′*^ values range between *−* 350 and 0.45 kJ/mol glucose (Figure 2b). Only one EFM had an overall stoichiometry with a positive Δ*G*^*m′*^, indicating that the reaction-reversibility constraints were set appropriately. In Figure 5, we consider the ATP/glucose yield of all net conversions as function of thermodynamic and reaction information. We highlighted particular net conversions in specific colors. (Important to note is that we use the term “Homoacetic” to refer to the exclusive production of acetate from glucose, yielding two acetate and two CO_2_ molecules (cf. (Zheng et al., 2014)) and not the homogeneous production of acetic acid from gaseous substrates like CO_2_/H_2_/CO (Karekar et al., 2022).)

**Figure 5:**
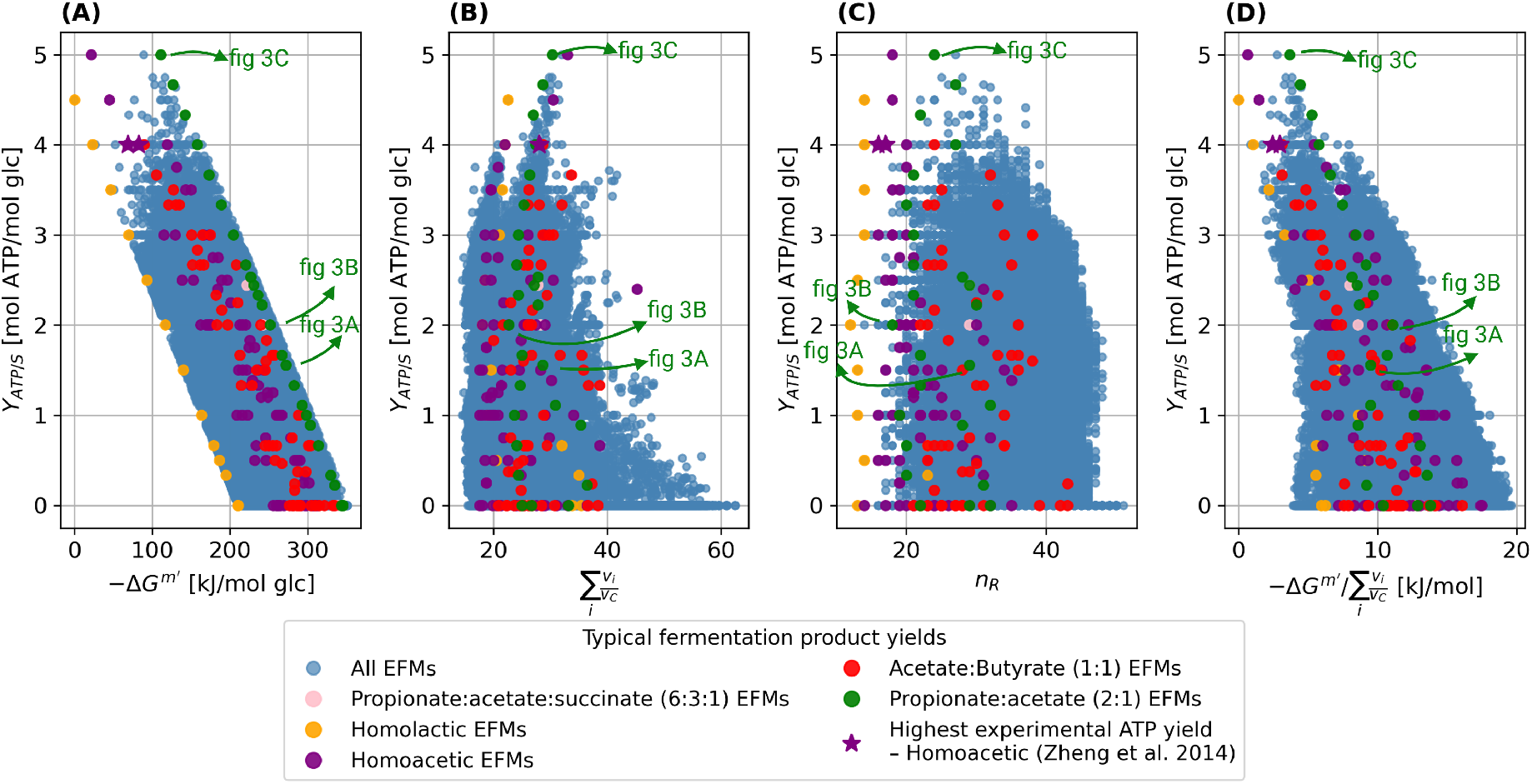
Thermodynamic assessment of all net conversions. ATP yield on glucose as a function of: (A). minus the standard Gibbs free energy change of the net conversion, 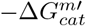, (B). the sum of all the flux ratio weights of reactions in the shortest EFM of the net conversion, (C). the number of reactions in the shortest EFM of the net conversion, and (D). the normalised standard Gibbs free energy, 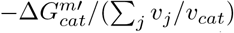. Some relevant catabolic product yields from literature are highlighted in a different color than blue. All the remaining net conversions are shown in blue. Note that “Homoacetic” in this work refers to the fermentation of glucose to two acetate and two CO_2_ molecules.

The Gibbs free energy change under standard conditions of a net conversion 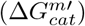 equals: i. the free energy released during the substrate-to-product conversion 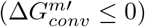 plus the free energy required for ATP synthesis 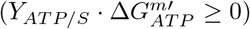 (Equation 1) and ii. the flux-ratio weighted sum of the standard free energies of the reactions of its corresponding EFM,

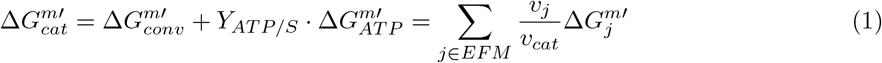

Each flux ratio corresponds to a reaction flux (*v*_*j*_; boxed numbers in the pathway figures 3 and 4) divided by the net conversion rate (*v*_*cat*_ equal to 1; since all net conversion are normalised per 1 glucose). In Figure 5, selected EFMs whose overall stoichiometry corresponds to those reported in Spormann (2023) and with the highest ATP yield on glucose experimentally measured (Zheng et al., 2014) are highlighted. Figure 5A displays the ATP yield on glucose for all conversions as function of their absolute standard Gibbs free energy change 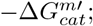 the standard driving force). It indicates that higher ATP yields correlate with lower driving forces as expected from equation 1. Net conversions with the same carbon conversion (all with the same 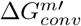 value) all fall on the same line in Figure 5A. For instance, homolactic fermentation of glucose (yellow markers) occurs with 12 different ATP yields on glucose, ranging from 0 to 4.5 ATP/glucose. These lines all have the same slope equal to 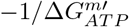. The intersection with the x-axis is the 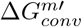 (because *Y*_*ATP/S*_=0). EFMs yielding lactate, or a 2:1 propionate-to-acetate ratio, vary only in their ATP yield, while EFMs producing acetate and a 1:1 butyrate-to-acetate ratio also differ in other products. For example, either H_2_ and CO_2_ or formate is produced. Because this alters the 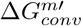, the datapoints for these stoichiometries do not fall on this line. This figure indicates that the fermentation of glucose into 1.33 propionate and 0.67 acetate has the most favourable driving force under the considered standard conditions and lactate fermentation the worst. We initially hypothesized that the 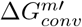 increases with the number of reactions: when glucose is broken down further, this would yield more driving force. However, this is not the case (Figure S3).

Figure 5B displays the ATP yield on glucose for all conversions as function of the sum of their flux-ratio weights. Equation 1 indicates that when those flux-ratio weights are high, the standard Gibbs free energy changes of the reactions tend to be low (indicating close to equilibrium operation of these reactions, requiring more enzyme for appreciable fluxes). This figure does not indicate any consistent correlation between ATP yield and the sum of their flux-ratio weights, neither across all net conversions and nor between those that have the same carbon conversion (i.e., the non-blue markers).

Figure 5C displays the ATP yield on glucose for all conversions as function of the number of reactions in their shortest EFM. Again no clear correlation is observed. Some observations include that the longest pathways have an ATP yield of 0 (Figure S4), that lactate fermentation corresponds to the shortest EFM, and that fermentation of glucose to equimolar amounts of acetate and butyrate tends to be among the longer pathways corresponding to known conversion profiles.

Finally, figure 5D displays the ATP yield on glucose for all conversions as function of the flux-ratio weight normalised driving force 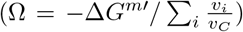. This x-axis measure equals 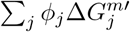 (with 0 *≤ ϕ*_*j*_ = *v*_*j*_*/* ∑_*i*_ *v*_*i*_ *≤* 1 and ∑_*j*_ *ϕ*_*j*_ = 1) and is high when the standard free energy changes of the reactions are each high (suggesting a lower enzyme cost (Noor et al., 2016)). The value of this x-axis is equal to the highest-possible value of max-minimal driving force (MDF) analysis (Noor et al., 2014) if no concentration bounds were supplied (Bruggeman et al., 2026). This makes it a rigorous estimate for thermodynamic favorability when Δ*G*^*m′*^ of individual reactions cannot be estimated. We found that no individual reaction is related to a favorable ATP yield nor Ω (Figure S5). This measure indicates that homoacetic and propionate/acetate fermentation are thermodynamically favorable per reaction and can also reach high ATP/glucose yields.

## 3 Discussion

This paper focuses on the microbial diversity of the anaerobic fermentation of glucose into catabolic products. Glucose is the main component of cellulose, one of the most abundant biopolymers on our planet (Bar-On et al., 2018), which makes anaerobic glucose fermentation (glucose catabolism) one of the most active processes. The aim of this work was to assess the diversity of microbial glucose catabolism in terms of its pathways, their thermodynamics, and yields of catabolic products and ATP. We computed those traits from a stoichiometric model of a pan-catabolic network of anaerobic glucose fermentation.

We compiled the pan-catabolic network from the literature. We acknowledge that it may be incomplete. We therefore provide all the Python scripts, which we used for this paper, to allow for either a more in depth or more focused analysis of anaerobic microbial fermentation, using our network as a template.

A catabolism is generally described in terms of its net conversion (Spormann, 2023), stating the yields of catabolic prodict and charged energy carriers on the energy source. We calculated the net conversions, by enumerating all feasible steady-state flux routes through our pan-catabolic network. In stoichiometric theory such routes are called elementary flux modes (EFMs) (Schuster et al., 2000), mathematical objects that can be computed with existing software (Terzer & Stelling, 2008). EFMs are metabolic pathways with a minimal number of reactions, of which none is redundant, with all reactions classified as (physiologically) irreversible operating in the corresponding direction.

Since many more EFMs generally exist than net conversion, we enumerated 60 million of them for them which gave rise to 195,629 unique conversions, of which about 50% had a non-zero ATP yield. We found 110,895 unique product stoichiometries. In the SI section S2, we explain how 60 million EFMs can arise via a combinatorial explosion of solutions. That many EFMs can give rise to the same net conversion also indicates that the catabolic network of a microbe cannot be predicted from its net conversion, but requires a catabolic network reconstruction from its sequenced genome.

We calculated many more net conversions than currently known from experimental studies. We doubt whether all of the computed conversions occur in nature. Many of them may be suboptimal for reasons not considered in this paper: kinetic, thermodynamic, ecological or otherwise. More in depth analysis of our results, using kinetic and thermodynamic methods such as max-min driving force (MDF) (Noor et al., 2014) or enzyme-cost minimisation (ECM) (Noor et al., 2016), may identify infeasibilities in some of these EFMs that were hidden from our methods. We refrained from such analysis here, because these methods require additional assumptions, of which some might be species or condition dependent (internal concentration bounds and kinetic parameters).

We view catabolic network reconstruction from sequenced genomes of anaerobic microorganisms as a promising next step to identify the catabolic reactions their genomes encode, to assess their potential for catalysis of specific net conversions. Performing such a computational analysis for a large set of species will indicate biases in the pathways responsible for anaerobic glucose fermentation.

We also found that the same conversion of glucose into fermentation products can give rise to varying ATP/glucose yields. This also indicates that the ATP yield cannot always be deduced from the stoichiometry of the fermentation products. This was already known for aerobically respiring microbes such as *Escherichia coli*, which have branched respiratory chains with different proton over electron stoichiometries (Poole & Cook, 2000).

Net conversions are often assessed on the basis of their Gibbs free energy change under standard conditions. Those with more negative values are then considered to be more favorable from a thermodynamic perspective, because less extreme substrate and product ratios are needed to drive those processes. Beause the 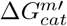 of a net conversion equals a flux-ratio weighted sum of the standard Δ*G*_*j*_ values of its reactions, considering only 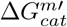 may obscure the comparison of pathways that differ greatly in the number of reactions and flux-ratio weights. Our analysis does not indicate any clear correlation of the ATP/glucose yield with those values. Thus, longer pathways do not necessarily lead to higher ATP yields. While the best thermodynamic estimates of pathway performance are most likely MDF and ECM, these measures depend on one’s access to kinetic information or one’s ability to make assumptions about ranges of feasible intracellular metabolite concentrations. The measure that we introduced 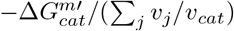 − does not require such inputs, and therefore estimates the distribution of thermodynamic driving forces with minimal ambiguity. In fact, 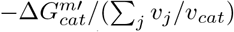 equals the MDF result when no concentration bounds are imposed (Bruggeman et al., 2026).

This work illustrates the potential of computational biology of microbial metabolism using mathematically-rigorous, computable concepts. It suggests that the metabolic potential of microbial species is currently greatly underestimated. We hope that the enhanced understanding of the diversity of microbial metabolism that this study offers benefits future exploitation of renewable resources.

## 4 Methods

The pipeline as presented in Bruggeman et al. (2026) was followed. Here, we give a short description of the method.

### 4.1 Model creation

The constraint-based modelling software COBRApy (Ebrahim et al., 2013) was used for verification and simulation. All enzyme-catalysed and transport reactions in the pan-catabolic network as presented in **Supplementary Text** were added to the model. The elemental balance of all internal reactions was verified using the elemental compositions of metabolites from the KEGG database (Kanehisa & Goto, 2000; Kanehisa et al., 2025). In COBRApy, the reactions are represented in a stoichiometric matrix (*N* ), where the stoichiometry of each reaction is represented as a column, with the stoichiometric coefficients in the row of the corresponding metabolites. Exchange reactions, in which an extracellular metabolite is consumed or produced without any counterpart, were added for glucose, the carbon products, H_2_,ATP, ADP, P_i_, H_2_O and H^+^. Directionality is controlled by the reaction bounds: a negative lower bound allows uptake, while setting the lower bound to zero restricts the reaction to secretion only. To verify whether all products could be made in this model, flux balance analysis (FBA) (Orth et al., 2010) was employed to optimise production of each product. Additionally, FBA was used to verify that ATP cannot be produced from ADP and P_i_ in the absence of glucose uptake.

### 4.2 Enumeration of elementary flux modes

A pathway can be formally defined by balancing all internal metabolites at steady state, such that knowledge of one flux determines all others. This is exactly the definition of an elementary flux mode (EFM) (Schuster & Hilgetag, 1994; Schuster et al., 2000). In terms of a stoichiometric model, this means that the flux vector *v* of an EFM must guarantee steady state of all metabolites (Eq. 2).

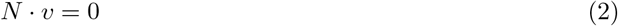

EFMs have two additional properties: they are non-decomposable, meaning no EFM is a linear combination of other EFMs, and they have exactly one degree of freedom, meaning that fixing one flux determines all others. We used the software package EFMTool (Terzer & Stelling, 2008) to enumerate all EFMs in the model. For simplicity, we normalised all EFMs to consume 1 glucose molecule. Then, we determined all unique overall stoichiometries, i.e. the EFMs that have a unique conversion of substrate(s) into product(s), with a unique ATP yield. For each unique overall stoichiometry, we chose the shortest EFM, i.e. the EFM with the fewest active reactions, to represent the unique overall stoichiometry. If two equally long EFMs led to the same overall stoichiometry, the first one was chosen.

### 4.3 Calculation of Δ*G*^*m′*^ values

We calculated the Δ*G*^*m′*^ of each overall stoichiometry using the eQuilibrator API (Noor et al., 2013), assuming standard physiological conditions (pH 7, 1 mM for all metabolites). For the calculation of the number of reactions (*n*_*R*_) and the sum of fluxes (∑_*j*_ *v*_*j*_*/v*_*cat*_), only enzyme-catalysed reactions were considered. The following reactions were thus excluded: exchange reactions and the ATP, ADP, and P_i_ transport reactions, as these are pseudoreactions; and the H_2_O and CO_2_ transport reactions, as these proceed by diffusion rather than enzyme catalysis.

## 5 Data Availability

The Python code that was used to generate the results of this work is available on https://github.com/SystemsBioinformatics/FermentationEFMs.

## 6 Acknowledgements

The authors would like to thank Dr Robbert Kleerebezem, Natalia Nadal Alemany and Wessel de Kok for their advice in assembling the pan-catabolic network. The figures were compiled in with BioRender.com. Maaike Remeijer and Frank Bruggeman received support from NWO-XL grant OCENW.XL21.XL21.007 ‘Taking Control of Metabolism in Microbial Cell Factories by Applying Noncanonical Redox Cofactors’. Christoff Odendaal received salary support from the European Union’s Horizon research and innovation programme under Grant Agreement No 10113540 (Nutritive). This work used the Dutch national e-infrastructure with the support of the SURF Cooperative using grant no. EINF-16049.

