## Supplementary information for "Comprehensive calculation of product and ATP yields for glucose fermentation"

### S1 Supplementary Text: Construction of pan-catabolic network

#### S1.1 General information

We constructed a non-exhaustive pan-catabolic network of typical anaerobic microbial glucose fermentation pathways. *Pan-catabolic* refers to the fact that we did not restrict our network to reactions co-occuring on the same genome. This document details the assumptions made and literature consulted based on which this network was constructed. The detailed model containing all reaction and metabolite identifiers, reaction equations, gene-protein-reaction associations, model construction comments, and metabolite chemical formulae is available in **Supplementary Table S1**. Note that some reactions in our model map onto multiple reactions in KEGG or to multiple EC numbers, proteins, or genes. On those cases, we provide all of the identifiers, genes, or proteins associated with the reaction as it appears in our model. Our model is divided into two comparments: the cytosol (c) and the extracellular environment (ex) from which substrates are taken up and to which products are exported.

As a point of departure, the chapter on *Fermentative Metabolism* by Spormann (2023) was used. We restricted our network to glucose fermentation pathways. The redox pairs $NADH/NAD^+$, $Fd^{2-}/Fd$ (reduced/oxidised ferredoxin), and $QH_2/Q$ (quinol/quinone) were internally balanced, as well as the consumption and production of free coenzyme A (CoA). Catabolism was modelled as ATP-generating: ATP was therefore net exported from the network, and $P_i$ and ADP net-imported. $H_2O$, $H^+$, and $CO_2$ could be freely consumed or exported.

NADPH is an electron donor to many anabolic reactions, while it is principally produced in the catabolism (Spaans et al. 2015; Chandel 2021). Unlike other redox pairs, therefore, it is not internally balanced. Including the possible net production of both ATP and NADPH would significantly increase the number of EFMs identified in our, we omitted $NADPH/NADP^+$-linked reactions from our model entirely.

Reactions were assigned to different subsystems to aid interpretability and human-readibility:

- Glycolysis (Embden-Meyerhof-Parnas pathway)
- PPP (pentose phosphate pathway - only the non-oxidative part)
- TCA Cycle (tricarboxylic acid cycle - only the reductive branch)
- Acetate
- Lactate
- Ethanol
- Butyrate
- Butanol
- Acetone
- Isopropanol
- Butanediol
- Propanoate (propionate)
- Electron carriers (hydrogenases, proton pumps and other enzymes that balance electrons)
- Transport (cross membrane)
- Exchange (import/export into model space)

To be maximally comprehensive, we compiled data from various databases as follows:

1. From KEGG (Kanehisa et al. 2025):
    - Reaction name
    - Metabolite name
    - Abbreviation of reaction or metabolite names
    - KEGG identifiers (for metabolites and reactions)
    - Chemical formula of reactions
    - Presence in *Clostridium pasteurianum* genome (specifically: *Clostridium pasteurianum* DSM 525 = ATCC 6013)
2. From UniProt ("UniProt: the universal protein knowledgebase in 2025" 2025):
    - EC Numbers
    - Reaction reversibility

- UniProt Entry Name
- UniProt ID
- Protein Name
- Gene name

3. From EBI-EMBL (Thakur et al. 2026):
   - Reaction reversibility (if not available on UniProt)
4. From ExPASy (Gasteiger et al. 2003):
   - Used to search for appropriate Reaction Names when KEGG name was too elaborate
   - Where EC numbers relate to multiple reaction, identify all related reactions using ExPASy
5. From eQuilibrator (Beber et al. 2022):
   - If no other information found, assume reversible unless the $\Delta G^{0\prime}$ provided on eQuilibrator is highly negative (< -20 kJ/mol).

Wherever there was still uncertainty about reaction reversibility, gene-protein-reaction relations, the ability of *C. pasteurianum* to catalyze given reaction, or anything else, we explicitly cited literature **Supplementary Table S1**. Below, we highlight these cases and reference the appropriate sources.

#### S1.2 Reversibility

For the following reactions, the reversibility was set based on direct literature sources:

- Phosphotransferase system (PTS), irreversible (Deutscher, Francke, and Postma 2006; Meadow et al. 2005)
- Phosphofructokinase (PFK), irreversible (Kuil, Nurminen, and Maris 2023)
- Fructose-6-phosphate phosphoketolase (F6PPK), irreversible (Hellgren et al. 2020)
- D-xylulose-5-phosphate phosphoketolase (X5PPK), irreversible (Hellgren et al. 2020)
- Pyruvate:ferredoxin oxidoreductase (PORf), revesible (Chen, Aman, et al. 2018)
- Pyruvate-formate lyase (PFL), reversible (Zelcbuch et al. 2016)
- Phosphate acetyltransferase (PTA), reversible (Miyake et al. 2000)
- Pyruvate carboxylase (PC), irreversible (Hanson and Owen 2013)
- $NAD^+$-dependent-malate dehydrogenase (MDHx), reversible (Berndsen and Bell 2024)
- Fumarase (FUM), reversible (Mescam, Vinnakota, and Beard 2011)
- $H^+$-translocating fumarate reductase (FR), reversible: proton-motive force (PMF)-forming direction describe by Spormann (2023), PMF-consuming direction described for *Bacillus sibtilis* by Schirawski and Unden (1998)
- Lactate:NAD+, ferredoxin oxidoreductase (LDHf), irreversible (Spormann 2023)
- For acetaldehyde oxidoreductase (ACALDORx), despite the reverse activity having been identified *in vitro* (White et al. 1989), the extreme thermodynamically forward nature of the reaction (-55.6 kJ/mol (Beber et al. 2022)) convinced us to model it as irreversible
- Dihydroxyacetone kinase (DHAK), irreversible (Monniot et al. 2012)
- Glycerol dehydrogenase (GDH), reversible (Leichus and Blanchard 1994)
- 1,3-propanediol dehydrogenase (PDOD), reversible (Malaoui and Marczak 2000)
- Acetoacetyl-CoA thiolase (THIO), reversible (Goudarzi 2019; Kursula et al. 2002)
- NADH-dependent 3-hydroxybutyryl-CoA dehydrogenase (HBDx), reversible (Liu et al. 2021)

- Electron-bifurcating butyryl-CoA dehydrogenase (BCDH_bifurc), reversible (Imran, Wiley, and Lubner 2024)
- Butyryl-CoA dehydrogenase (NAD-dependent) (BDHx), irreversible (Hu et al. 2013)
- Bifunctional Butanal-butanol dehydorgenase (AADH_BUTO), reversible (Pony et al. 2020)
- Acetoacetate decarboxylase (ACACDC), irreversible (Neece and Fridovich 1967)
- Isopropanol Dehydrogenase (NAD(+)-dependent) (IPADHx), reversible in its NADPH-coupled form (Ismaiel et al. 1993; Molinas et al. 2003), assume the same for the NADH-coupled form
- Lactoyl-CoA dehydratase (LACOAD), reversible (Hofmeister and Wolfgang Buckel 1992)
- NAD+-dependent acryloyl-CoA reductase (PROPCOAORx), irreversible (Hetzel et al. 2003)
- Formate dehydrogenase (FDH), reversible (Scherer and Rudolf K. Thauer 1978)

Assume succinate dehydrogenase (SDH) is irreversible, despite it typically being reversible ("UniProt: the universal protein knowledgebase in 2025" 2025). This decision is made because modelling SDH as reversible leads to ATP-forming cycles and the direction typically described in literature is forward.

Also assume that alpha-ketoglutarate ferredoxin oxidoreductase (AKGDHf) is irreversible to prevent the occurence of a loop which can convert reduced ferredoxin (FDred) to NADH. Chen, Li, et al. (2019) make it clear, however, that this reaction is biochemically reversible.

*Clostridium thermocellum* has a highly reversible NADH-coupled lactate dehydrogenase (LDHx) (Özkan et al. 2004). Assume irreversible, as we do not have a lactate consuming model. Similarly, quinone-dependent D-lactate dehydrogenase (LDHq) is reported in the context of lactate consumption in *Priopionibacterium freudenreichii* (Spormann 2023). Assume irreversible to prevent NADH conversion fo $QH_2$, which could yield a PMF and, consequently, ATP.

Butyrate kinase (BUTK), which is known to be reversible (Hartmanis 1987), is modelled as being irreversible to prevent ATP-dissipating loops.

Hydrogenase (HYDf) is reversible (Vignais, Billoud, and Meyer 2001), but we assume it is irreversible to present ATP-producing loops in combination with RNF. An NADH-coupled hydrogenase (HYDx) was identified in *Thermoanaerobacter tengcongensis*, only functions at low hydrogen partial pressure (Soboh, Linder, and Hedderich 2004). Assume it is irreversible like HYDf.

Assume that NADH-quinone oxidoreductase (NUO), detected in *Schwenella oneidensis* (Madsen and TerAvest 2019), is reversible like the mitochondrial NADH oxidoreductase ("UniProt: the universal protein knowledgebase in 2025" 2025).

##### S1.3 Detected in *C. pasteurianum*

Reactions which were ascertained as being possible in *C. pasteurianum* based in direct literature sources:

- PC, present (Sabra et al. 2016)
- MDHx, present (Pyne et al. 2016)
- SDH, indicated as absent from *C. pasteurianum* (Pyne et al. 2016), despite being predicted as being present on UniProt
- Succinate CoA synthetase (SCS) indicated as absent from *C. pasteurianum* (Pyne et al. 2016), despite being predicted as being present on UniProt
- The presence of aconitase (ACO) in *C. pasteurianum* is strongly suggested by the presence of icocitrate dehydrogenase (IDH) (Pyne et al. 2016)
- Malolactic enzyme (MALCL), present (Pyne et al. 2016)
- Bifunctional acetaldehyde-ethanol dehydorgenase (AADH_ETH), present (Poehlein et al. 2015)

- BDHx, present (Spormann 2023)
- AADH_BUTO, present (Poehlein et al. 2015)
- Butyrate-acetoacetate CoA-transferase (ACBUTCOAT), present (Pyne et al. 2016)
- Acetate-acetoacetate CoA-transferase (ACACCOAT), present (Pyne et al. 2016)
- ACACDC, present (Pyne et al. 2016)
- FDH, present (Rudolf K. Thauer, Fuchs, and Käufer 1975)
- ATP synthase (ATPase), present (Das and Ljungdahl 2003)

#### S1.4 Other modelling decisions

In terms of stoichiometry, all of the ferredoxins in the model were considered to be 2[4Fe-4S]-ferredoxin - capable of transferring two electrons at a time. 2[4Fe-4S]-ferredoxins are more typical of central carbon metabolism while [2Fe-2S]-ferredoxins, which transfer only one electron at a time, are more typical of nitrogen fixation reactios (Vallières et al. 2024; Sahin et al. 2025). This is not exclusively true, but was the operative assumption of this model.

The lactate:NAD+,ferredoxin oxidoreductase (LDHf), an electon-bifurcating lactate dehydrogenase was reported in *Acetobacterium woodii* (Weghoff, Bertsch, and Müller 2015) and hypothesisded in *Clostridium tyrobutyricum* (Munier et al. 2023). In these cases, LDHf is reported in the context of lactate-consumption. Spormann (2023), however, reports it in the context of pyruvate reduction to lactate. In the latter case, the primary literature is not cited. To include this reaction, but to prevent the formation of NADH-oxidising loops, we only include the reaction in the direction of lactate formation.

Lactate-malate transhydrogenase (MALTR) is included, but it bears noting that is is only known in *Micrococcus lactilyticus* (Allen 1966).

Although a reverse instance of the AADH_ETH was observed by Pony et al. (2020), we model it as irreversible as our model is not ethanol-consuming and modelling it as reversible leads to the occurence of NADH-oxidising loops. Likewise, as ethanol is not a substrate in the model, Acetaldehyde oxidoreductase (ALDH_ETH) is assumed irreversible.

It seems like, besides alcohol dehydrogenase (adhE), which is a two-step channeled reaction going from acetyl-CoA to ethanol, and butyryl-CoA to butanol, there is an NADPH-dependent butanol dehydrogenase (bdh) which catalyses the reaction from butyraldehyde to butanol (Yoo et al. 2015). Assume, therefore, that butanal and acetaldehyde can also be released from the NADH-dependent alcohol dehydrogenase.

1-propanol dehydrogenase (PODH) is hypothetical, but it is also the enzyme responsible for the last step in producing propane-1,3-diol from glycerol, the 3-hydroxypropionaldehyde reductas. Its presence is inferred from teh fact that production of propane-1-ol is known in some propionic acid bacteria (Walther and François 2016).

Although UniProt indicate that HBDx has NADP/NADPH as electron acceptor/donor pair, Jungermann et al. (1973) show that the pathway from acetyl-CoA to butyrate must be NADH-consuming and not NADPH-consuming. This is also consistent with the way the pathway is represented by W. Buckel and R. K. Thauer (2013) and Pyne et al. (2016).

The PMF-forming stoichiometries of RNF and the energy-converting hydrogenase (ECH), as well as that of ATPase, were assumed the same as in Schoelmerich et al. (2020), W. Buckel and R. K. Thauer (2013), and Katsyv and Müller (2022).

HydABC - an electron-confurcating hydrogenase - performs the same function as another enzyme HndABCD (Kpebe et al. 2018). We model the reaction only once and name it HydABC.

Otto et al. (1980) showed that the export of lactate can generate a proton gradient which was later shown by Von Ballmoos, Wiedenmann, and Dimroth (2009) to be usable for ATP production via ATP synthase. Wolfgang Buckel (2021) reviews this as well.

### S2 Supplementary Text S2: Combinatorial explosion

The 60 million EFMs we computed arise from a combinatorial explosion. To illustrate this, we use a method developed by Kelk et al. (2012).

For the production of propionate, acetate and succinate (in a 6:3:1 stoichiometry) with an ATP yield of 2 ATP/glucose, there were 8 unique EFMs (Figure S1A). Using the block diagonalized correlation matrix of fluxes of the reactions with variation in the flux yielded three subnetworks. In the first subnetwork (Figure S1A1), pyruvate is converted into acetyl-CoA, and the same net conversion can occur in two ways. In the second subnetwork, succinyl-CoA was converted into succinate and ATP was produced, either in one reaction step (SCS), or using 6 reaction steps (Figure S1A2). In this case, the biologically relevant option is the 1-step conversion, as likely less enzyme is required for the same net conversion flux. The third subnetwork had three options, all using the same sum of fluxes for the conversion of lactate, propionyl-CoA and acetyl-CoA into lactoyl-CoA, propionate and acetate (Figure S1A3). Stoichiometrically and thermodynamically, these three subnetworks are equivalent. Together, these three subnetworks cause $2 \cdot 2 \cdot 3 = 8$ options for the conversion of glucose into propionate, acetate and succinate with an ATP yield of 2 ATP/glucose.

For the production of propionate and acetate with a 2:1 ratio and an ATP yield of 5 (as shown in Figure 3C), there were 184 EFMs. Three subnetworks were determined, two of which with two pathway options and one with 46 options (the shortest network is shown in Figure 3C). In that case, it already becomes more difficult to determine the exact pathways of the options. In the case of the same conversion, but a lower ATP yield of 2 ATP/glucose (three of the networks are shown in Figure 4), there were 2,596 EFMs. However, no subnetworks are present (Figure S1C). This is caused by the covariance of each reaction with each other.

Here, we showed that the combinatorics of the networks are complex. For a small subset of EFMs, the combinations are disentangled readily. However, with increasing network size, the flux values have too much covariance.

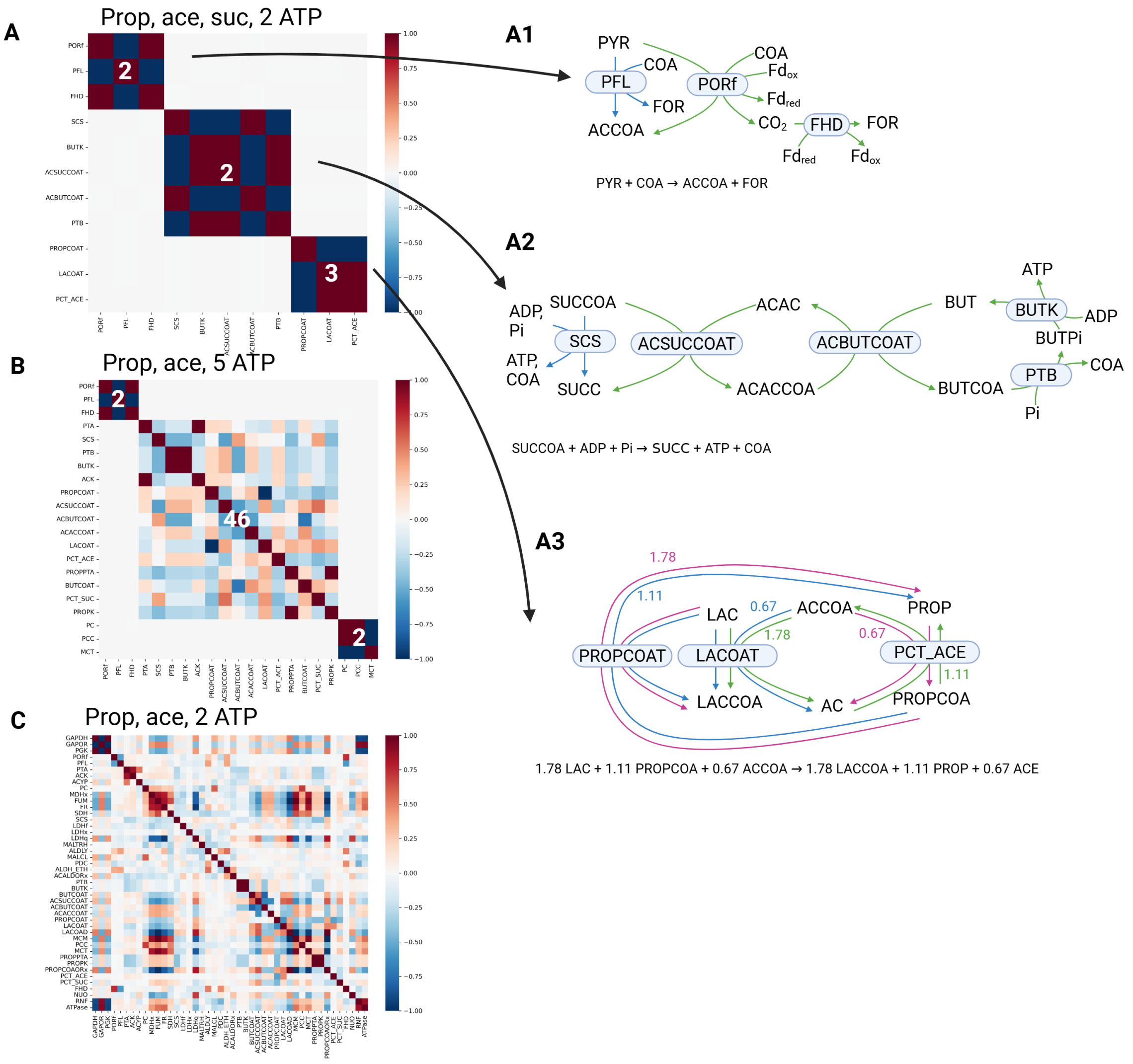


Figure S1: **Combinatorial explosion in small subnetworks can be explained using block diagonalized correlation matrices of varying fluxes.** (A) 8 EFMs were found for the production of propionate, acetate and succinate (with a 6:3:1 ratio) and an ATP yield of 2 ATP/glucose. Three subnetworks were found, (A1) for the conversion of pyruvate into acetyl-CoA and formate, (A2) for the conversion of succinyl-CoA into succinate, with production of ATP and (A3) for the conversion of lactate, propionyl-CoA and acetyl-CoA into lactoyl-CoA, propionate and acetate. (B) 184 EFMs were found for the production of propionate and acetate (with a 2:1 ratio) with an ATP yield of 5 ATP/glucose. These EFMs are subdivided into three subnetworks, two with 2 pathway options and one with 46 pathway options. (C). 2,596 EFMs were found for the production of propionate and acetate (with a 2:1 ratio) with an ATP yield of 2 ATP/glucose. No further distinction into subnetworks was found.

#### S3 Supplementary Figures

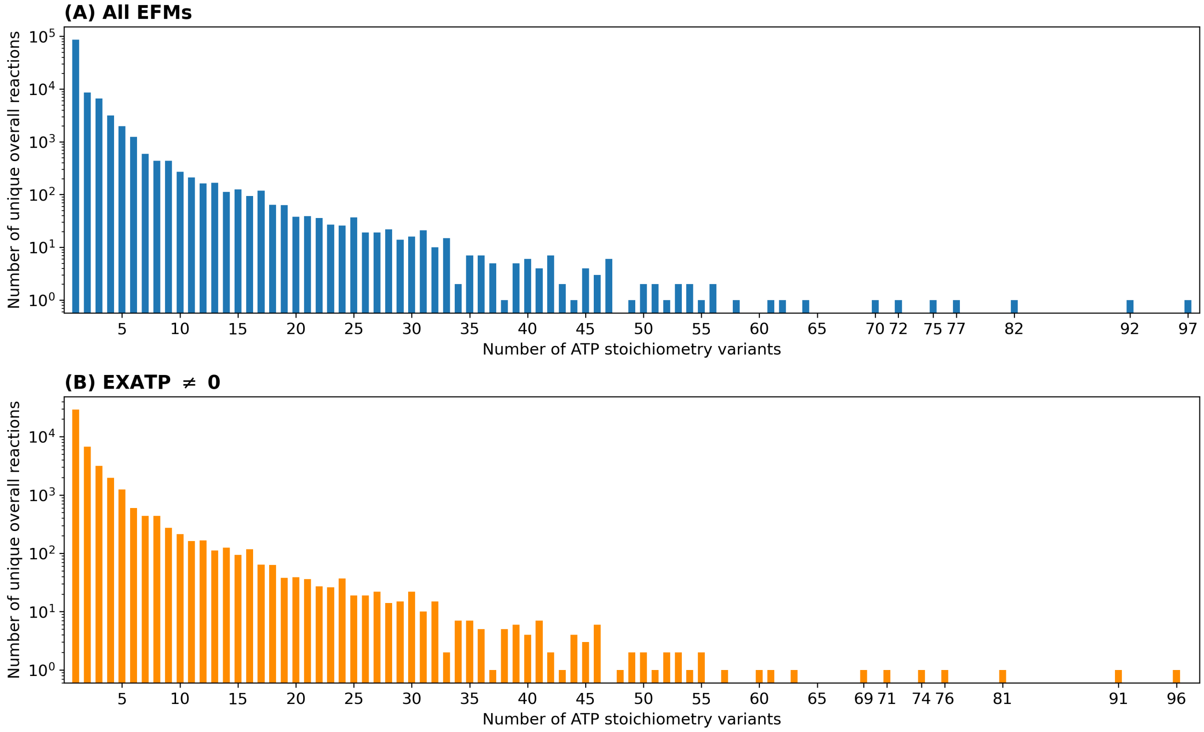


Figure S2: Counts of unique overall reactions that have a given ATP/glucose yield fo (A) all EFMs and (B) EFMs with non-zero ATP yields.

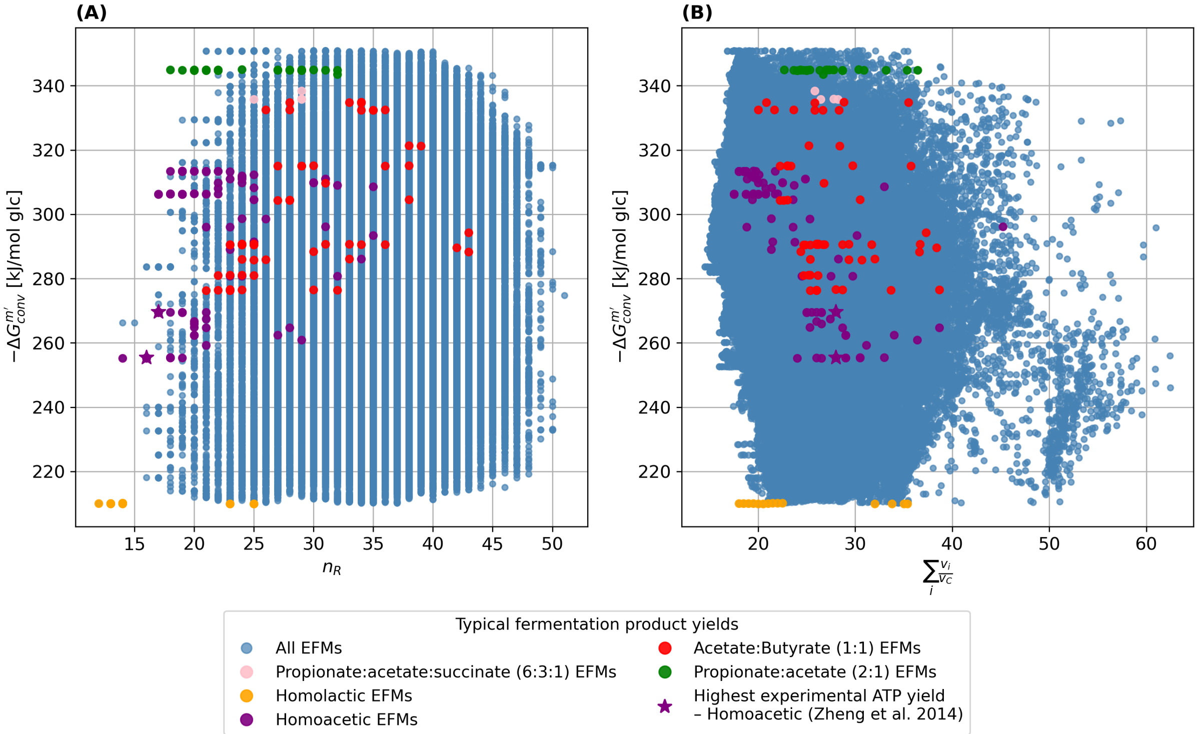


Figure S3: $-\Delta G_{conv}^{m\prime}$ as a function of the number of reaction steps ($n_R$) (A) and the sum of fluxes ($\sum_i \frac{v_i}{v_C}$) (B), for all EFMs (blue) and for selected common fermentation product ratios (red, purple, green, orange, pink; see legend). The purple star indicates the EFM matching the highest experimentally reported ATP yield for homoacetic fermentation (Zheng et al., 2014). No clear relationship is apparent between the thermodynamic driving force of the conversion and either the number of reaction steps or the total flux required by the pathway.

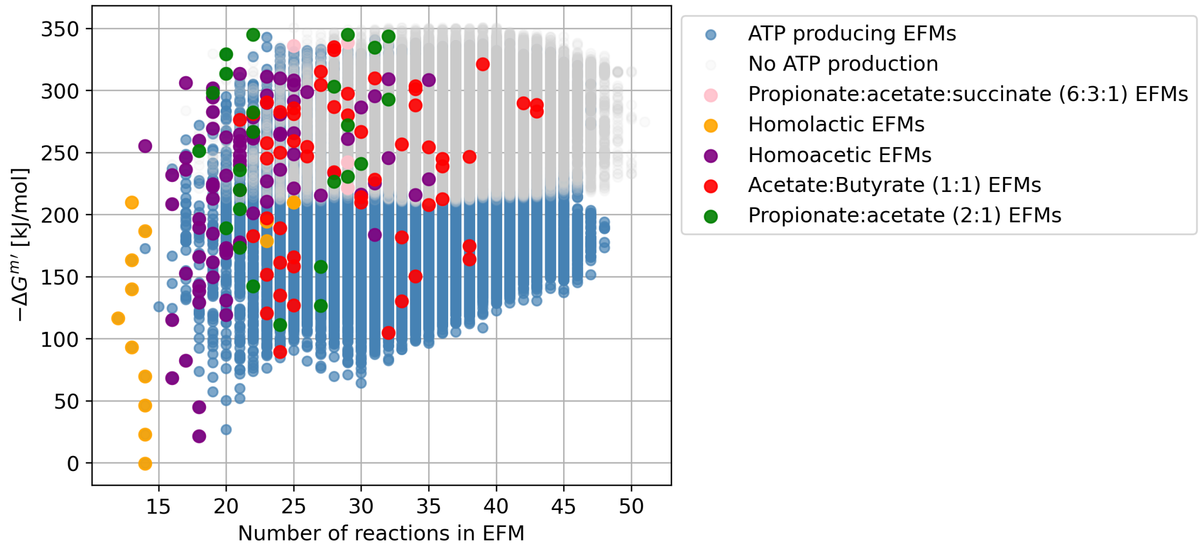


Figure S4: $-\Delta G^{m\prime}$ as a function of the number of reactions in the EFM ($n_R$), for all EFMs (blue), EFMs with zero ATP yield (grey), and selected common fermentation product ratios (red, purple, green, orange, pink; see legend). The EFMs with the highest number of reactions appear to be ones without ATP production.

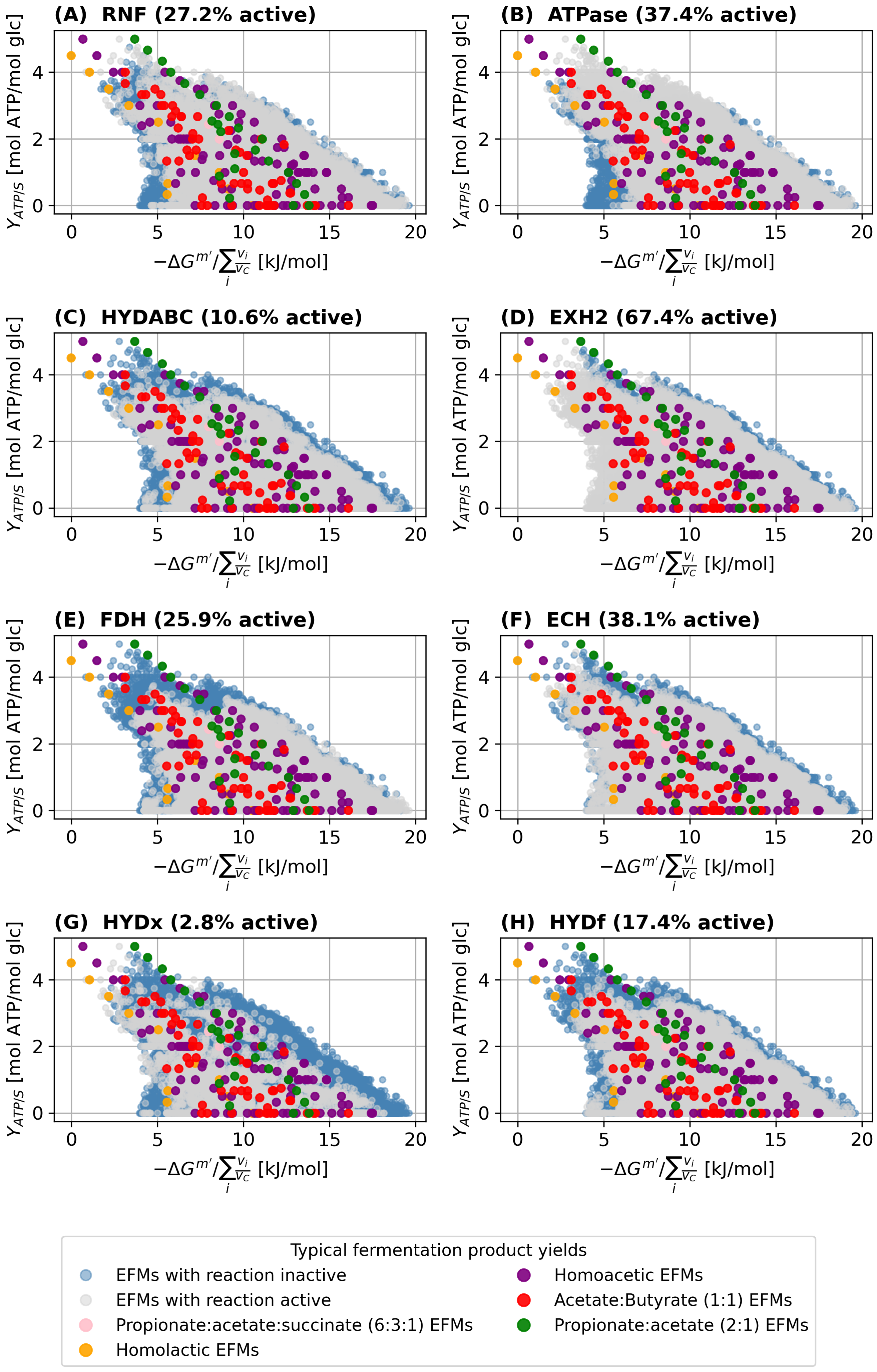
(A) RNF (27.2% active)
(B) ATPase (37.4% active)
(C) HYDABC (10.6% active)
(D) EXH2 (67.4% active)
(E) FDH (25.9% active)
(F) ECH (38.1% active)
(G) HYDx (2.8% active)
(H) HYDf (17.4% active)
$Y_{ATP/S}$ [mol ATP/mol glc]
$-\Delta G^{m'} / \sum_i \frac{v_i}{v_C}$ [kJ/mol]
0
2
4
0
5
10
15
20
Typical fermentation product yields
EFMs with reaction inactive
EFMs with reaction active
Propionate:acetate:succinate (6:3:1) EFMs
Homolactic EFMs
Homoacetic EFMs
Acetate:Butyrate (1:1) EFMs
Propionate:acetate (2:1) EFMs

Figure S5: Utilization of reactions involved in hydrogen production and ATP generation across EFMs. For each reaction, $Y_{\mathrm{ATP/S}}$ is plotted against $-\Delta G^{m\prime}/\sum_i \frac{v_i}{v_C}$, with EFMs in which the reaction is active shown in grey and those in which it is inactive shown in blue. Selected common fermentation product ratios are highlighted (red, purple, green, orange, pink; see legend). The percentage of EFMs in which the reaction is active is indicated in each panel title. Reactions shown: (A) ferredoxin:NAD$^+$ oxidoreductase (RNF), (B) ATP synthase (ATPase), (C) electron-bifurcating hydrogenase (HYDABC), (D) hydrogen exchange (EXH2), (E) formate dehydrogenase (FHD), (F) energy-converting hydrogenase (ECH), (G) NADH-dependent hydrogenase (HYDx), and (H) ferredoxin-dependent hydrogenase (HYDf).
